# Multi-epitope Based Peptide Vaccine Design Using Three Structural Proteins (S, E, and M) of SARS-CoV-2: An *In Silico* Approach

**DOI:** 10.1101/2020.06.13.149880

**Authors:** Arpita Singha Roy, Mahafujul Islam Quadery Tonmoy, Atqiya Fariha, Ithmam Hami, Ibrahim Khalil Afif, Md. Adnan Munim, Mohammad Rahanur Alam, Md. Shahadat Hossain

## Abstract

Severe Acute Respiratory Syndrome Coronavirus-2 (SARS-CoV-2) is the novel coronavirus responsible for the ongoing pandemic of coronavirus disease (COVID-19). No sustainable treatment option is available so far to tackle such a public health threat. Therefore, designing a suitable vaccine to overcome this hurdle asks for immediate attention. In this study, we targeted for a design of multi-epitope based vaccine using immunoinformatics tools. We considered the structural proteins S, E and, M of SARS-CoV-2, since they facilitate the infection of the virus into host cell and using different bioinformatics tools and servers, we predicted multiple B-cell and T-cell epitopes having potential for the required vaccine design. Phylogenetic analysis provided insight on ancestral molecular changes and molecular evolutionary relationship of S, E, and M proteins. Based on the antigenicity and surface accessibility of these proteins, eight epitopes were selected by various B cell and T cell epitope prediction tools. Molecular docking was executed to interpret the binding interactions of these epitopes and three potential epitopes WTAGAAAYY, YVYSRVKNL, and GTITVEELK were selected for their noticeable higher binding affinity scores −9.1, −7.4, and −7.0 kcal/mol, respectively. Targeted epitopes had 91.09% population coverage worldwide. In summary, we identified three epitopes having the most significant properties of designing the peptide-based vaccine against SARS-CoV-2.

## Background

History suggests that humanity is always challenged by emerging viruses and viral infections in terms of livelihood and economic progress in a population. The current situation of ongoing pandemic of corona virus disease (COVID 19) verily supports that history. The disease came to recognition by the World Health Organization (WHO) as a pandemic on 11^th^ March, 2020 and has caused a global emergency across 210 countries and territories around the world and 2 international conveyances [1]. As of today, 11^th^ June, 2020 at 18.09 (GMT +6), a confirmed report of 3,295,032 active infected cases with 2% criticality and 421,363 deaths has been found. (https://www.worldometers.info/coronavirus/). But the disease originated back in December, 2019 in Wuhan City, Hubei Province, China in the form of cluster of pneumonia like symptoms which then quickly transcended the border to spread across the globe within a short span of time [2].

The causative agent behind the infection was first designated as 2019 novel coronavirus (2019-nCoV) by the World Health Organization (WHO)[3]. The novel coronavirus was further renamed to severe acute respiratory syndrome coronovirus-2 or SARS-CoV-2 because of its genomic similarity of 79.5% and 96% at nucleotide level, respectively with SARS-CoV and bat coronavirus [4, 5].Through phylogenetic analysis, SARS-CoV-2 has been categorized under the family Coronaviridae and order Nidoviralae and has shown an origin in bat as a natural host [6, 7]. The SARS-CoV-2 is similar to the SARS-CoV and MERS-CoV of the same genus *Betacoronavirus* within the same family in terms of infectivity in humans. The latter two viruses caused pandemic situations as well in recent years and were the reasons for thousands of death across the world [8–10]. As for the structural similarity, the SARS-CoV-2 contains a positive single stranded RNA as its genetic element having a genomic length of around 30 kilobases [11]. The encoded proteins by this genome are structural and non-structural in kind and the major structural proteins are spike (S) glycoprotein, membrane (M) protein, envelope (E) protein, and nucleocapsid (N) protein [12–14]. Especially the exposed parts of these proteins contain domains necessary for infection into the host cells and account for antigenicity. The similarity between SARS-CoV and SARS-CoV-2 extends to the structure of spike glycoprotein which may be exploited for a potential vaccine design because the S protein has two major subunits namely S1 and S2 which assist during viral infection into the host cell. S1 subunit contains receptor binding domain (RBD) and N-terminal domain (NTD) where RBD binds to the receptor of the host cell known as angiotensin converting enzyme-2 (ACE2) [15, 16]. The envelope (E) protein also has its role in establishing a series of involvement from pathogenesis to viral assembly by not only interacting with the host cell protein but also maintains a defining connection with all the other structural proteins (M, S, N proteins) of the virus [17]. Moreover, the exposed portion of M protein outside the membrane provides us an opportunity to design a suitable epitope-based subunit vaccine.

Currently, different therapeutic strategies are being utilized by the scientific community to combat this dangerous spread of COVID-19, most of which are opted to develop novel drugs or vaccines. Chinese traditional drugs named *ShuFengJieDu* capsules and *LianHua Qing Wen* capsules had been applied to some of the preliminary cases before they were reported to be effective. But no clinical trials to this date have ever proven them to be safe enough[18]. Some other drugs such as Remdesivir and Chloroquine had also been reported to be effective against COVID-19 through in-vitro trials. However, no authentic clinical trial has justified such a claim so far [18, 19].Besides, SARS-CoV-2 is dispersing too fast across almost all the countries of the world and is mutating in an unbelievable manner. Due to the high mutation rate in the genome of SARS-CoV-2 single epitope will not suffice to provide for a successful vaccine; rather, a multiple epitope based vaccine may do the trick here[20].

At the current stage of the pandemic, it will be highly insensible to develop a vaccine through classical approach *in vitro* that involves identification, isolation and culture of pathogenic viruses. Following this particular process will be too expensive and time consuming which is not desirable at all given the circumstances. A very sustainable way to overcome such hurdles would be to design peptide vaccine by genome and proteome analysis of the virus using computational methods. Since the genome and proteome sequencing of SARS-CoV-2 has already been done, it is only rational that we take control measures by making the best out of computation-based analysis to design therapeutic targets. In this study, we explored the S, E, and M proteins of SARS-CoV-2 by using different *in silico* tools and servers to predict B-Cell and T-Cell epitopes to eventually design an effective epitope-based vaccine. The predicted epitopes were analyzed further to check their antigenicity and surface accessibility. Epitope-allele interaction was investigated through molecular docking. A phylogenetic analysis was undertaken to identify molecular evolutionary relationship of the selected S, E and M proteins. The study was concluded with the introduction of a properly designed vaccine from the most suitable of epitopes.

## Materials and Methods

### Retrieval of Protein Sequences

The FASTA format of S, E, and M protein sequences of SARS-CoV-2 from various geographical areas: Australia, China, USA, Finland, India, Sweden, South Korea were retrieved from National Center for Biotechnology Information (NCBI) (https://www.ncbi.nlm.nih.gov/). Then, the BLAST program from NCBI was used to derive the similar sequences against the proteins.

### Phylogenetic tree construction and analysis

The S, E, and M protein sequences of SARS-CoV-2, SARS-CoV, MERS and common human coronavirus strains (229E, NL63, OC43, and HKU1) were targeted for this phylogenetic study. All the sequences of the proteins retrieved from the NCBI BLASTP result were aligned separately through the ClustalW algorithm by utilizing the MEGA (version 10.0.5) [21].All the required parameters for the alignment analysis were used as default program in the software. The aligned sequences were then visualized with the Jalview (version 2.11.0) [22]for the observation of consensus and conserved sequences.. The phylogenetic trees were built using the neighbor-joining tree function and default analysis preferences in MEGA.

### Membrane Topology Analysis

Epitopes of a protein must be in the exposed regions to mount sufficient immune response. The membrane topology of these proteins was analyzed using TMHMM v2.0 server[23]and was later cross-referenced with the InterPro server[24, 25]. The outer membrane regions of these proteins were selected for further analysis.

### Antigenicity prediction

A vaccine candidate must elicit sufficient antigenic response as antigenicity of epitopes play a crucial role to provoke adequate immune response. VaxiJen v2.0 server[26]calculates antigenicity depending on physicochemical properties of proteins with the threshold value 0.4 (for viral protein sequence).

### B cell epitope Identification

B cell epitopes presented on the virus surface proteins are recognized by B lymphocytes to elicit immune response. Based on artificial neural network, ABCpred v2.0 server[27–30] predicted the linear B cell epitopes. The epitopes were cross-referenced with Immune Epitope Database (IEDB)[31], which uses amino acid scales and Hidden Markov Models (HMM) as prediction method. Moreover, Kolaskar and TongaonkarAngenicity, Parker Hydrophilicity Prediction tools from IEDB were explored to determine the antigenicity and hydrophilicity properties of these selected epitopes.

### T cell epitope Identification

T cell epitopesconsist of a group of amino acids, presented by antigen presenting cell (APC) in the bound form with major histocompatibility (MHC) molecules to mount T cell mediated immune response. Prediction of T cell epitopes was performed by using NetCTL tool, which utilizes MHC binding affinity, proteosomal processing, TAP transport[32–34]. The tool predicts half-maximal inhibitory concentration (IC50) values of epitopes based on Artificial Neural Network (ANN)[35, 36].The lengths for epitopes were set at 9.0 and 15.0 for MHC I molecule and MHC II molecular, respectively.

### Molecular Docking Analysis of HLA and Epitopes

The three-dimensional structure of targeted T-cell epitopes were modeled by the PEP-FOLD3server [37],a peptide structure predicting tool. The best 3D structure generated by this server was selected as a ligand for docking analysis. Molecular docking was performed by the virtual screening tool Pyrx[38]through its Autodock vina [39, 40] program in order to analyze the interactions among our proposed epitopes and different HLA molecules. Protein Data bank provided most of the pdb files of HLA molecules for docking study whereas Phyre2 protein prediction server (Protein Homology/analogy Recognition Engine v2.0) [41] was used to generate the 3D models of some HLA molecules whose pdb structureswere not available in PDB. Discovery Studio(v4.5) [42]prepared these HLA molecules as macromolecules by removing water, non-polar hydrogen and unnecessary molecules.Pdbfiles were converted into pdbqt files and the default grid box parameters were maintained with exhaustiveness value 8.The binding interactions of epitope-HLA molecules were visualized using UCSF Chimera 1.13rc[43].

### Population coverage prediction

Determination of Population coverage for individual is essential as epitopes may exhibit variety in their binding sites during interacting with different HLA allele. The IEDB population coverage calculation tool [44]was utilized to determine the percentage of people expected to respond to a specific number of MHC-restricted epitopes around the world.

## Result & Discussion

One of the most prolific ways to reduce the load of viral diseases is to design novel vaccines [45–48]. In terms of SARS-CoV-2, continuous research projects are being held to figure out the biological characteristics, genomics and proteomics and pathophysiology of the virus around the world [49]. Immunoinformatics tools have always been a medium to develop multi-epitope based vaccines for some of the dangerous viral diseases such as rhinovirus[50], dengue virus[51],chikungunya virus[52] etc. Consistent with that idea, this study aimed at providing for a peptide based vaccine design against SARS-CoV-2 using different bioinformatics tools by exploring structural proteins of virus namely spike (S) glycoprotein, membrane (M) protein and envelope (E) protein[53, 54]responsible for virulence mechanism and pathogenic pathways. A total of 13 sequences of S protein **(Figure S4)**, 13 sequences of M protein **(Figure S5)**, and 6 sequences of E protein (**Figure S6)** were selected for analysis.

S, E, and M protein sequences from different strains of MERS, SARS-CoV, SARS-CoV-2 (**Table S1&S3)** and common human coronavirus, (**Table S2**) were aligned with the phylogenetic tree construction using the built-in inClustalW algorithm in the MEGA (version 10.0.5).

Phylogenetic trees represented the ancestral molecular changes, evolutionary history and the molecular evolutionary relationship on account of S, M, and E proteins, respectively. From the phylogenetic analysis of spike glycoprotein from the different strains of MERS, SARS-CoV SARS-CoV-2 and common human coronaviruses (Human coronavirus strains: 229E, NL63, OC43 and HKU1), we noticed that their ancestral and evolutionary relationship has been comprehensive and found relatable **(Figure S1)**. Analysis also showed that both human coronavirus OC43, and HKU1 are closely related to MERS virus whereas distantly related to the SARS and SARS-CoV-2. On the other hand, human coronavirus NL63, and 229E are closely related to each other. Phylogenetic analysis of the membrane proteins **(Figure S2)** of different strains of MERS, SARS-CoV, SARS-CoV-2 and envelop proteins **(Figure S3)** from diverse strains of MERS, SARS-CoV, SARS-Cov-2 along with common human coronaviruses strains (229E, NL63, OC43, and HKU1) showed that SARS-CoV and SARS-CoV-2 are closely related to MERS virus whereas OC43, and HKU1 strains are closely related, and also comparatively closer to the SARS-CoV, SARS-CoV-2, and MERS virus rather than the other closely related 229E and NL63 strains. Unlikely the ancestral relationship in regard of S and M proteins, analysis showed a different result for E protein that SARS-CoV and SARS-CoV-2 are closely related to MERS virus whereas human coronavirus 229E and NL63 are closely related, and also related to comparatively closer to the SARS-Cov, SARS-Cov-2 and MERS rather than the other closely related strain human coronavirus OC43 and HKU1.

Viral infection is facilitated by the outer portion of the domain of viral protein which binds with the receptor proteins of the host cell. So exposed portions of S, E, and M proteins of SARS-CoV-2 normally offer a large scope to induce antibody production against itself[55, 56].Therefore, it is necessary to design a therapeutic target based on those exposed portions of the virus. In order to do so, Interpro and TMHMM servers were used to predict the outer portions of each of the protein. The length of exposed regions (Non-cytoplasmic region) ranging from 1-1213, **(Figure 1)** (1-11, 35-75) and (1-19, 74-78) respectively for S, E, and M proteins were taken for our assessment. A vaccine candidate must have antigenic features to maximize its reliability to be an agent of immune response. To ensure the efficacy of the vaccine, antigenicity of the target sequences must be checked as it is crucial for the design of peptide-based vaccine. VaxiJen 2.0 server was used to predict antigenicity of all the respective proteins. The S protein region had an antigenic value of 0.4646 whereas values for the region of E, and M protein were 0.7282, and 0.5102, respectively.

Potential B cell epitopes play significant role by providing protection against viral diseases[57].Therefore, distinctive analysis techniques were utilized in this method for the prediction of a linear B cell epitope. Primary sequences of S, E, and M protein were checked through ABCpred and IEDB server to foresee B-cell epitopes. A total of 59, 5, and 19 B-cell epitopes for S, E, and M were predicted individually by ABCpred. From all anticipated epitopes, only 4 epitopes; S (2 epitopes), E (1 epitope), and M (1 epitope) presented on the outside of S, E, and M proteins were chosen (Table 1) with higher antigenicity score.Kolaskar and Tongaonkar antigenicity estimation tool **(Figure2)** was utilized to predict this antigenicity, score appeared in **Table1** and TMHMM server was used to check the surface accessibility.

**Table 1.** Predicted B-cell linear epitopes with ABCpred score, antigenicity and hydrophilicity score.

| Protein | Epitope | Position | ABCpred score | Antigenicity (IEDB) | Hydrophilicity (IEDB) |
| --- | --- | --- | --- | --- | --- |
| **S** | HRSYLTPGDSSSGWTA | 245-260 | 0.92 | 0.993 | 2.95 |
|  | CFTNVYADSFVIRGDE | 391-406 | 0.85 | 1.048 | 1.513 |
| **E** | NVSLVKPSFYVYSRVK | 48-63 | 0.8 | 1.113 | 0.45 |
| **M** | TITVEELKKLLE | 7-18 | 0.75 | 1.043 | 0.492 |

Based on ABCped, among these four identified epitopes of S protein, ‘HRSYLTPGDSSSGWTA’, and ‘CFTNVYADSFVIRGDE’ showed most elevated antigenic score of 0.92, and 0.85, respectively. Moreover, epitope ‘HRSYLTPGDSSSGWTA’ is positioned within the NTD region whereas ‘CFTNVYADSFVIRGDE’ is inside the RBD region. The E protein epitope ‘NVSLVKPSFYVYSRVK’ showed 0.8 antigenic score. This epitope is situated at 48 position of E protein. While at 7 position of M protein the ‘TITVEELKKLLE’ revealed 0.75 antigenic score and selected for M protein **(Table 1)**. According to IEDB, the ‘TITVEELKKLLE’ indicated most noteworthy antigenicity of 1.113 followed by the ‘CFTNVYADSFVIRGDE’, ‘TITVEELKKLLE’, and ‘HRSYLTPGDSSSGWTA’ with the antigenicity of 1.048, 1.043, and 0.993, respectively. The ABCpred score and VaxiJen antigencity results revealed the ability of all predicted peptides to expand barrier reactions within the host during SARS-CoV-2 infection as an extracellular part of transmembrane-protein. To discover the hydrophilicity of anticipated B-cell epitopes parker-hydrophilicity strategy was performed **(Figure3)**. With the hydrophilicity analysis, the ‘HRSYLTPGDSSSGWTA’ was found to have remarkable hydrophilicity with the value of 2.95, whereas hydrophilicity values of ‘CFTNVYADSFVIRGDE’, ‘TITVEELKKLLE’, and ‘NVSLVKPSFYVYSRVK’ were 1.513, 0.492, and 0.45, respectively **(Table 1)**.

B-cell epitope based vaccines have been quite popular for a long time; however, vaccines based on T-cell epitopes are currently in the trend as the CD8+ T cells produce a long term memory response in the host against the infected cell [58].That’s why prediction of T-cell epitopes is an unquestionable requirement for a vaccine to have preventive capacity. Some of the recent studies on vaccine development against COVID-19 shed light on only the T-cell epitopes whereas in our study, we have approached both eligible B and T cell epitopes of SARS-CoV-2 for a maximized yield in our vaccine design project[58].Here, IEDB server was utilized to evaluate the best T-Cell epitopes from the chosen protein sequences of S, E, and M protein. Moreover, antigenicity testing and screening of peptides were done with help of VaxiJen 2.0 server. In light of the high combinatorial score, the 4 best epitopes for MHC Class-I **(Table 2)** were chosen for additional investigation. For MHC class I assessment, NetMHCcons 1.1 Server is a consensus approach that combines the three best NetMHC, NetMHCpan, and PickPocket class strategy to provide the most accurate predictions. The MHC-I alleles for which the epitopes demonstrated higher affinity (IC50, 500nM) were chosen **(Table 2)**.

**Table 2.**
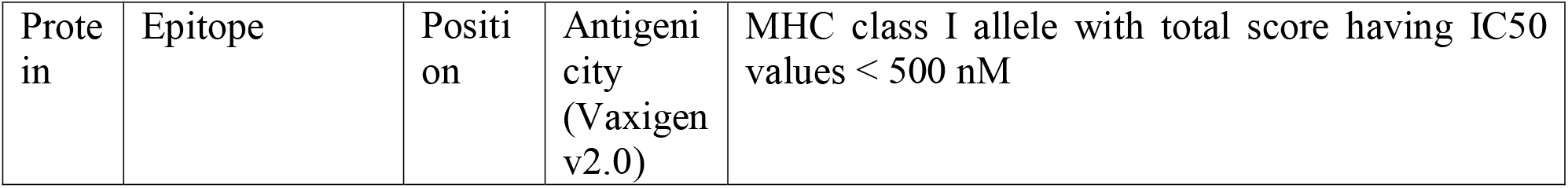
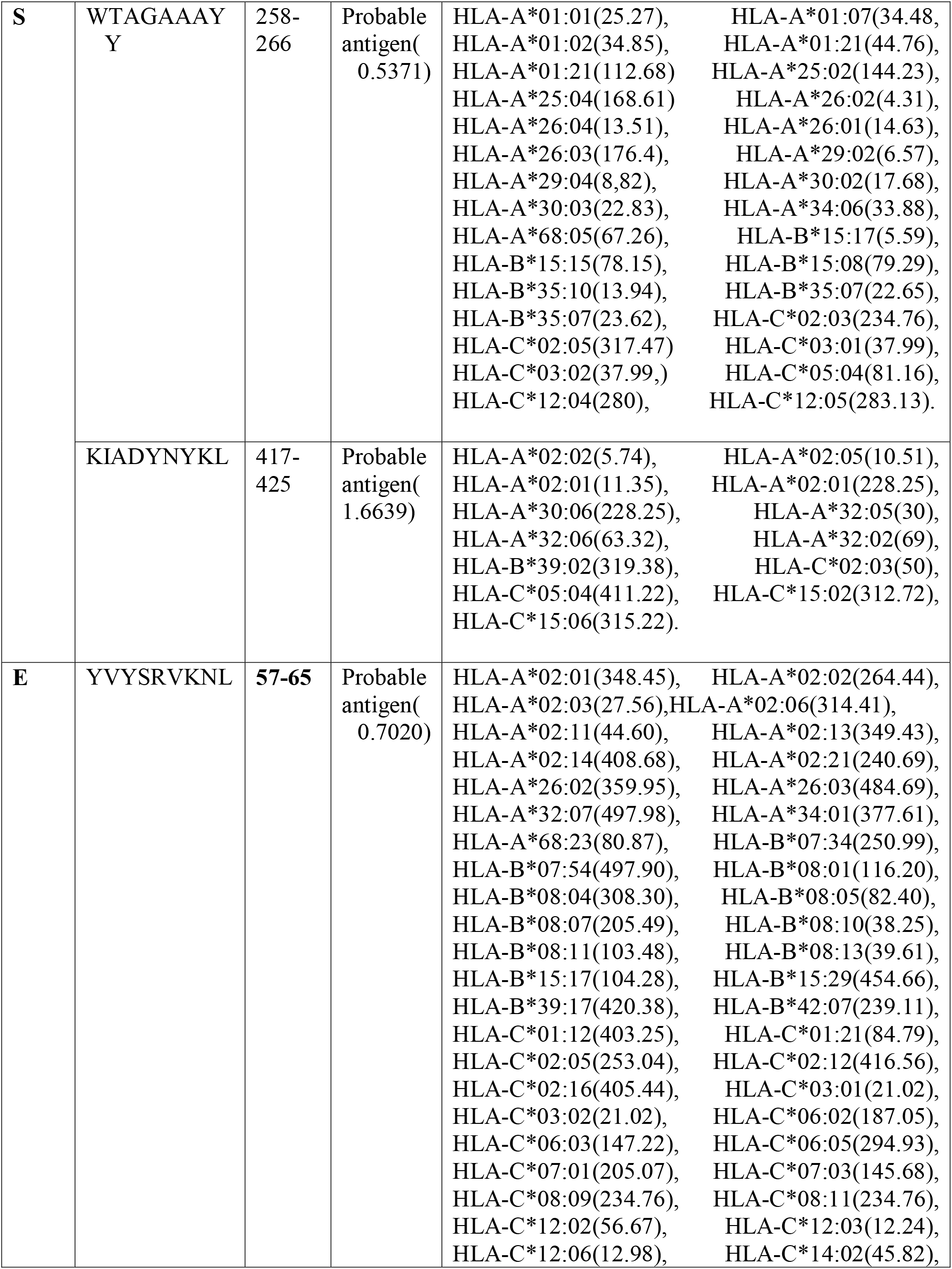
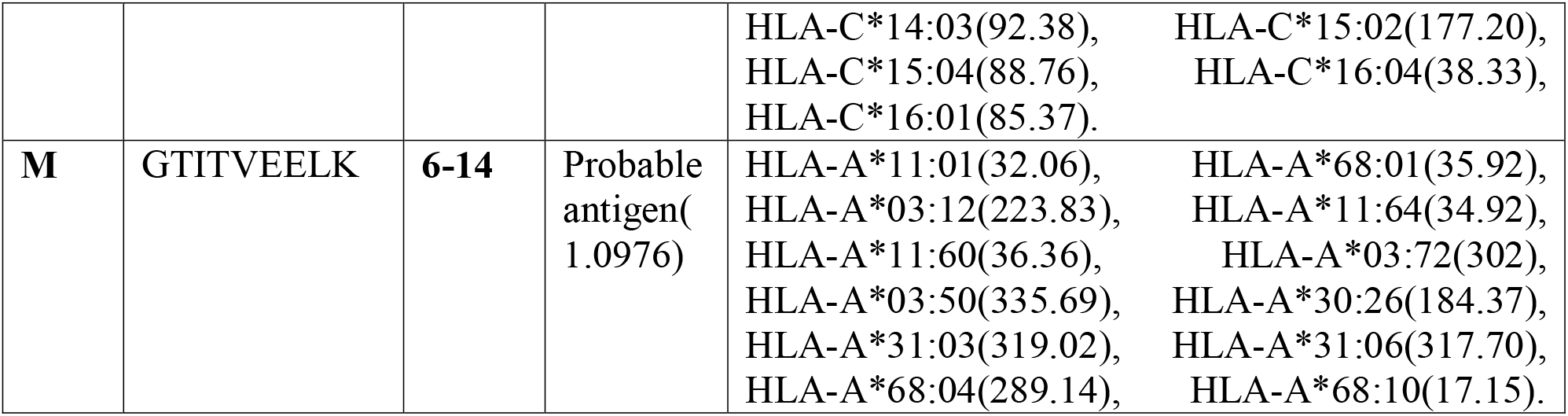
Predicted epitopes for CD8+ T-cell along with their interacting MHC class I alleles with affinity < 500 nM.

Between MHC class-I anticipated epitopes, a 9 mer epitope, ‘KIADYNYKL’ that is inside the RBD region of SARS-CoV-2 showed higher antigenicity score of 1.6639 followed by the ‘GTITVEELK’, ‘YVYSRVKNL’ and ‘WTAGAAAYY’ (within the NTD region of SARS-CoV-2) with antigenicity score of 1.0976, 0.7020, and 0.5371,respectively. For MHC Class-II, on the basis of high combinatorial score, the 4best epitopes **(Table S4)** were chosen for further analysis. NetMHCII 2.3 server was used to anticipate MHC-II binding prediction with HLA-DR, HLA-DQ and HLA-DP MHC class II alleles. The MHC-II alleles for which the epitopes demonstrated higher affinity (IC50, 500nM) were selected **(Table S4)**. The peptide ‘GVLTESNKKFLPFQQ’ that is within the RBD region of SARS-CoV-2 was viewed as increasingly antigenic for its higher antigenicity score 0.8200 followed by the ‘YFKIYSKHTPINLVR’ (inside the NTD area of SARS-CoV-2), ‘FYVYSRVKNLNSSRV’, ‘MADSNGTITVEELKK’ with antigenicity score of 0.8197, 0.6103, and 0.4367, individually.

Moreover, molecular docking was performed for the analysis of the binding affinity of our targeted eight epitopes from membrane, spike and enveloped protein with different HLA molecules. T-cell Epitope sequence with their docking score are showed in the **(Table 3)**. Among these eight selected T cell epitopes, ‘WTAGAAAYY’ bound in the binding pocket of HLA-B*35:01 allele (PDB ID:4PRN) showed the highest binding score of −9.1 kcal/mol **(Figure4& 5)**. Interaction between the epitope and the binding pocket atoms of HLA molecule showed in the **Figure 6**. On the other hand, ‘YVYSRVKNL’ bound with HLA-A02:03(PDB ID: 3OX8) and ‘GTITVEELK’ bound with HLA-A*11:01 (PDB ID: 6JOZ) with the binding affinity of −7.4, and −7 kcal/mol, respectively exhibit good binding interaction of these epitopes with HLA molecules. Another essential prerequisite in epitope based vaccine design is to determine population coverage as MHC polymorphism leads to the expression of different forms of HLA at considerably different rates among different ethnicities[59, 60]. On the basis of binding interaction of our targeted epitopes with their respective HLA alleles we finalized three epitopes for vaccine design having the highest binding affinity. Population coverage for these three epitopes demonstrated the consequences of the host genetic variety on the binding specificity of these targeted epitopes to class I HLA alleles. Their cumulative population coverage around the world 91.09% which revealed that these epitopes can cover about 91% **(Figure 7)** population from different regions of the world.

**Table 3.** Binding affinity of T-cell epitopes and HLA molecules.

| Protein name | Epitope seq. | Allele | Docking score(kcal/mol) |
| --- | --- | --- | --- |
| M Protein | GTITVEELK (MHC I) | HLA-A*11:01 (32.06) | -7 |
|  | MADSNGTITVEELKK (MHC II) | HLA-DQA1*01:02/DQB1*06:02 (215.20) | -5.5 |
| E Protein | YVYSRVKNL (MHC I) | HLA-A02:03(27.56) | -7.4 |
|  | FYVYSRVKNLNSSRV (MHC II) | HLA-DRB1*04:01 (12.50) | -6.2 |
| S Protein | WTAGAAAYY (MHC I) | HLA-B*35:01(23.62) | -9.1 |
|  | KIADYNNKL (MHC I) | HLA-A*02:01(11.35) | -6.1 |
|  | YFKIYSKHTPINLVR (MHC II) | HLA-DRB1*04:01(185.8) | -6.4 |
|  | GVLTESNKKFLPFQQ (MHC II) | HLA-DRB1*03:01(188.1) | -4.6 |

Most of the recent studies emphasized on only the spike protein of the virus given its wide range of conservancy across the viral population. But our work was dedicated to specific epitopes and domains of all the mentioned proteins (S, M, and E proteins) for an *in-silico* design of multi-epitope vaccine. Vaccine development against SARS-CoV-2 shed light on only the T-cell epitopes in some recent studies whereas in our study, we have approached both eligible B and T cell epitopes of SARS-CoV-2 for a maximized yield in our vaccine design project [61]. All the ideal criteria of a potential epitope had been pulled off by our selected epitopes of S, M, and E proteins.

## Conclusion

Different vaccine development strategies against SARS-CoV-2 are being opted along with ongoing trials on novel drug design. In our current study, we suggested three epitopes from conserved regions of spike, membrane and envelope proteins of SARS-CoV-2 as they have abilities to produce antigenic response. Their potentialities as vaccine candidate assured by their antigenic properties and strong binding affinity for MHC molecules. This *in-silico* study requires experimental validation to get cost effective epitope based peptide vaccine with higher efficacy against SARS-CoV-2.

## Acknowledgements

The authors acknowledge the Department of Biotechnology and Genetic Engineering, Noakhali Science and Technology University for providing the research facilities.

## Author Contributions

MSH conceived and designed this study ASR, MIQT, AF, IBKA, IH, MAM performed the experiment and analyzed the data. MSH, RA, ASR, IH wrote the manuscript. All authors reviewed the manuscript.

## Additional Information

## Supplementary Information

Supplementary information for this study may be found online in the Supplementary Information section.

---

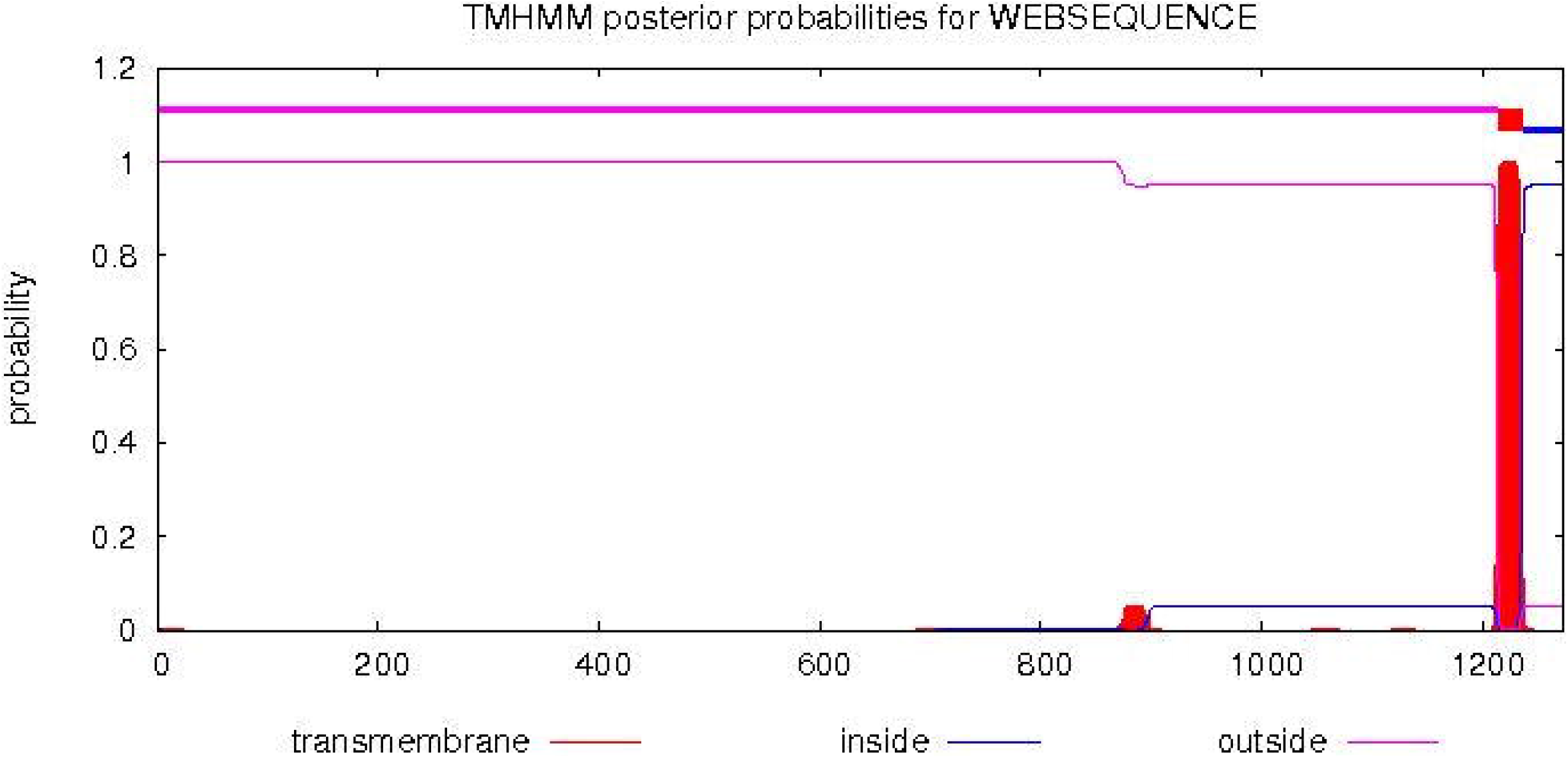

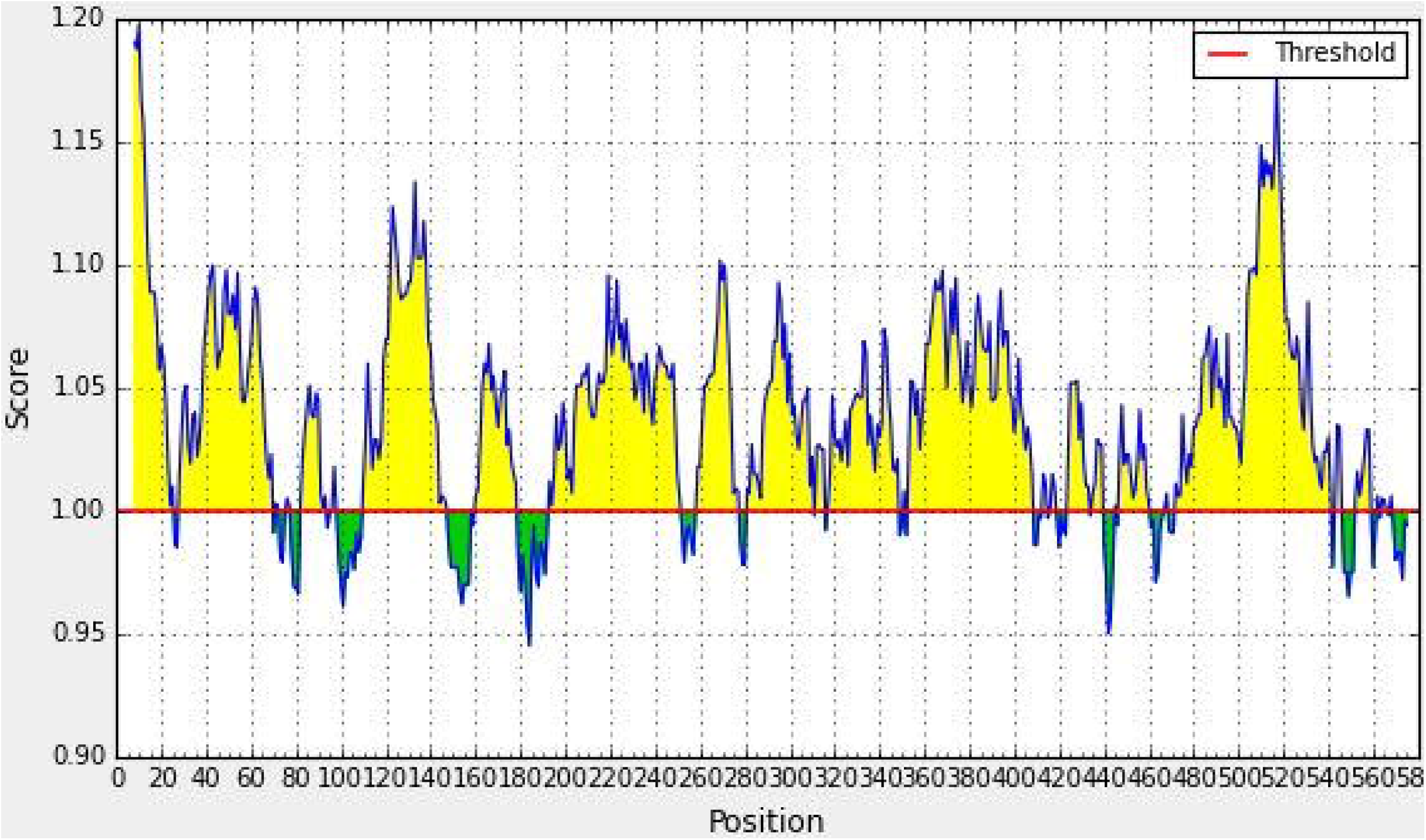

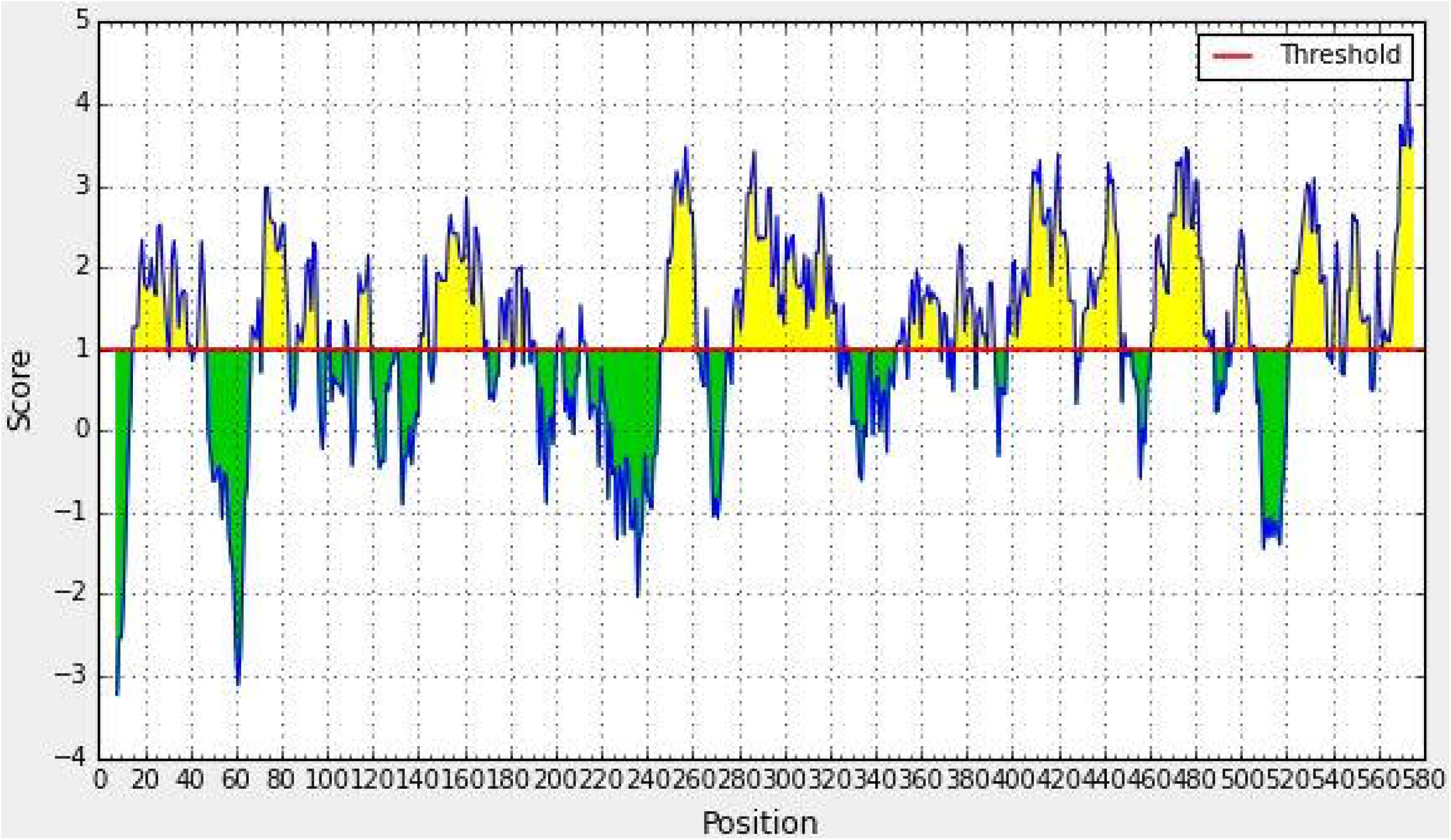

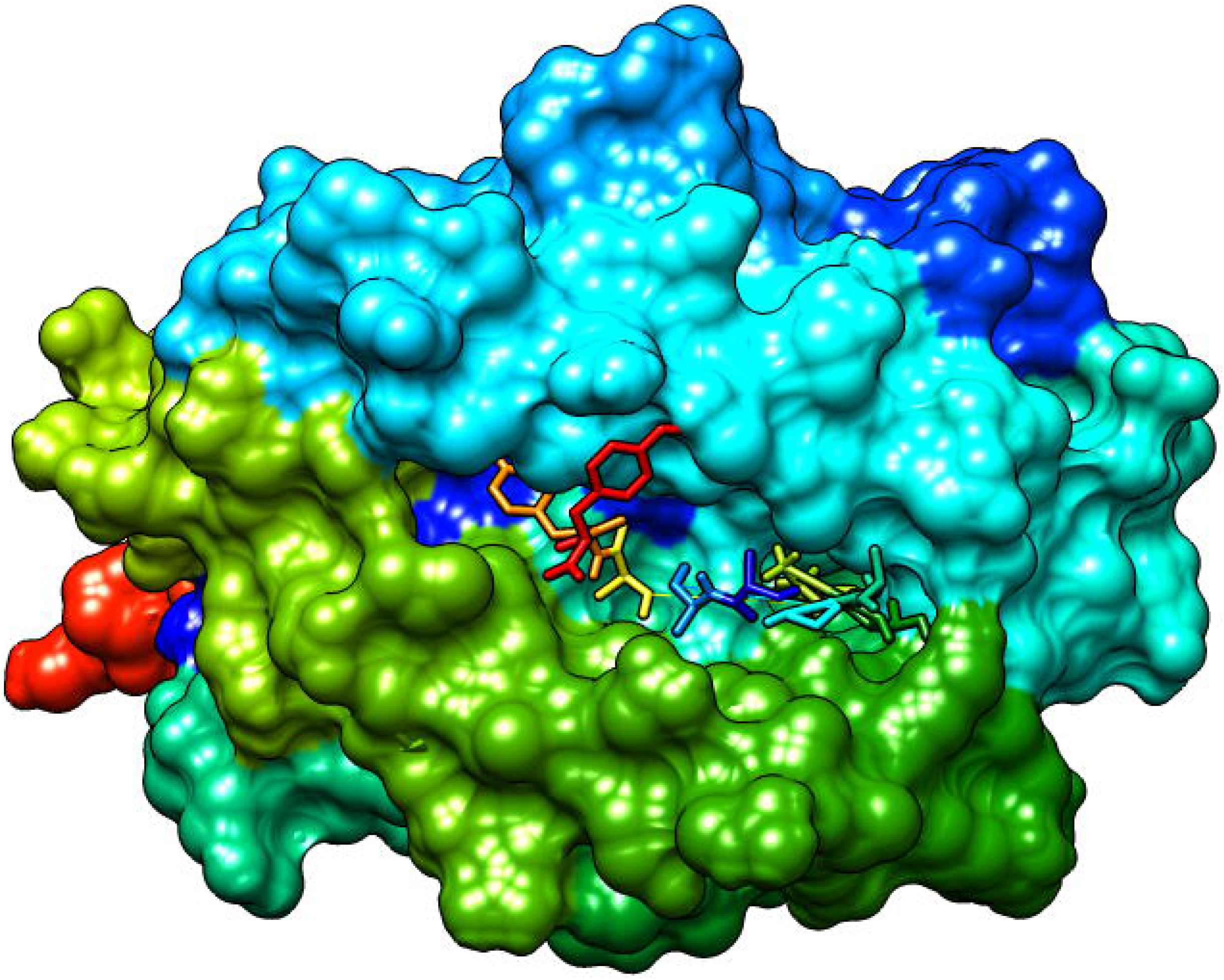

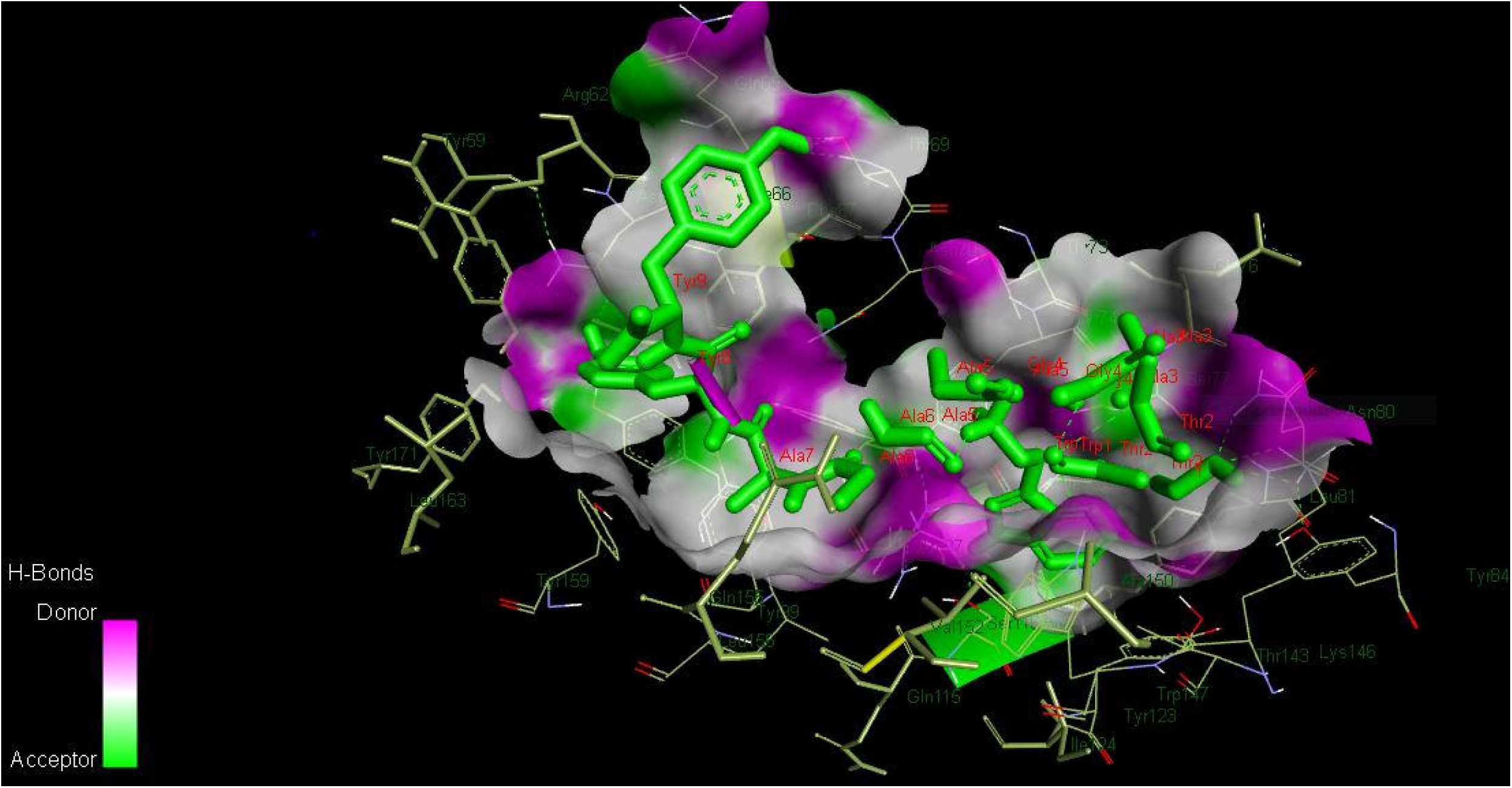

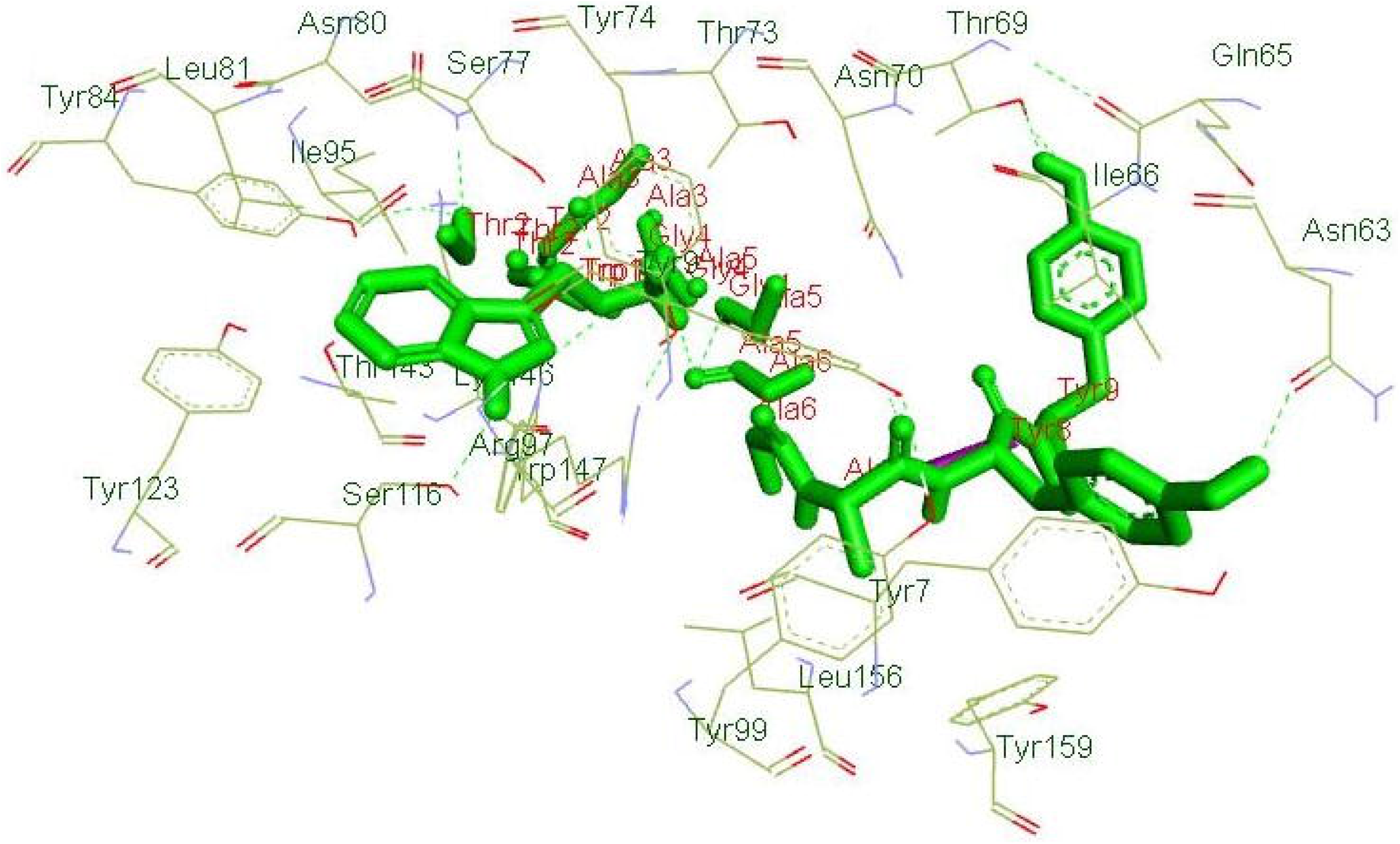

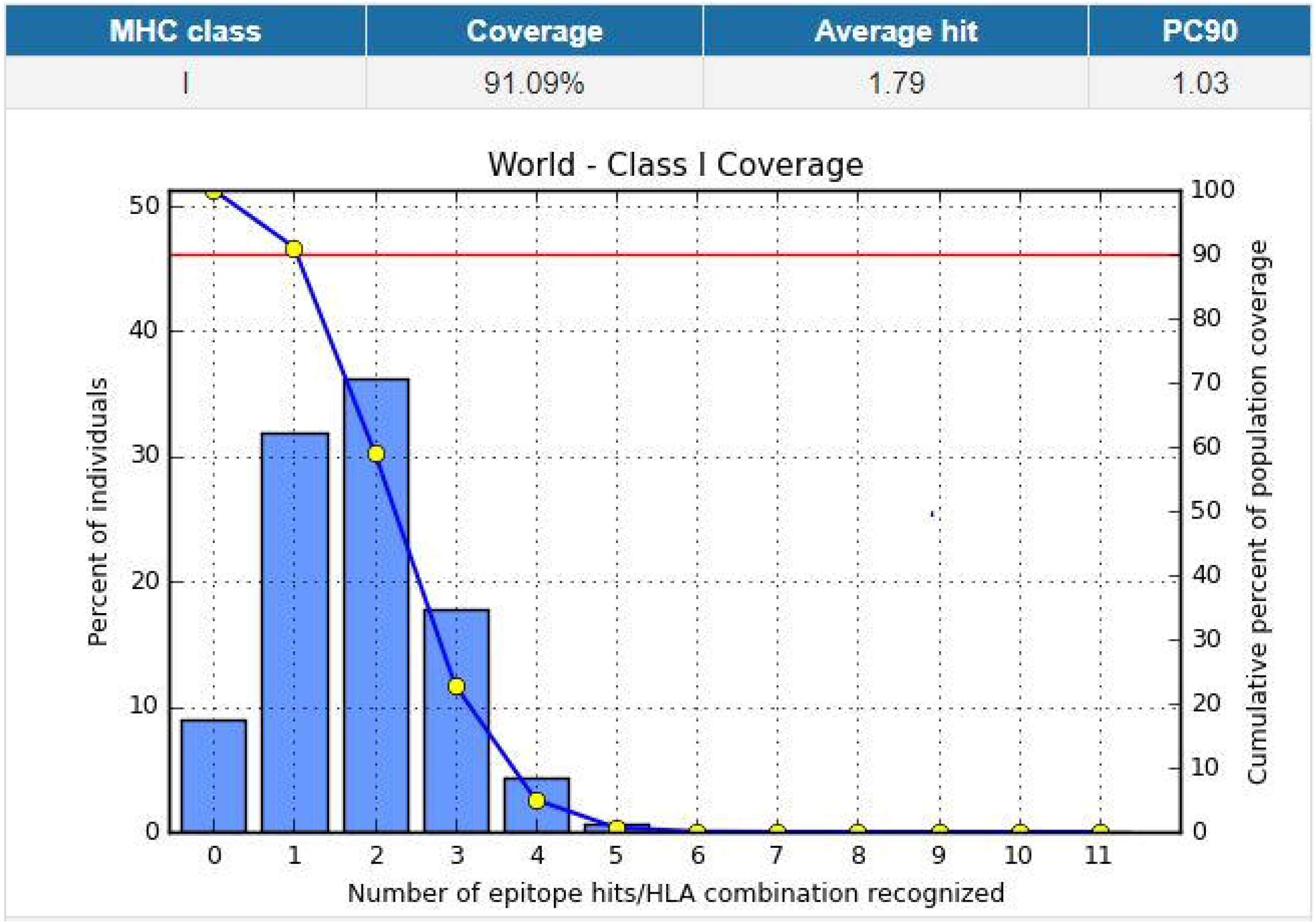

## References

1. Cucinotta, D. and M. Vanelli, WHO Declares COVID-19 a Pandemic. Acta Biomed, 2020. 91(1): p. 157–160.

2. Tang, P.F., et al., Expert consensus on management principles of orthopedic emergency in the epidemic of coronavirus disease 2019. Chin Med J (Engl), 2020. 133(9): p. 1096–1098.

3. Benvenuto, D., et al., The 2019-new coronavirus epidemic: Evidence for virus evolution. J Med Virol, 2020. 92(4): p. 455–459.

4. Zhou, P., et al., A pneumonia outbreak associated with a new coronavirus of probable bat origin. Nature, 2020. 579(7798): p. 270–273.

5. Zhu, N., et al., A Novel Coronavirus from Patients with Pneumonia in China, 2019. N Engl J Med, 2020. 382(8): p. 727–733.

6. Hui, D.S., et al., The continuing 2019-nCoV epidemic threat of novel coronaviruses to global health – The latest 2019 novel coronavirus outbreak in Wuhan, China. Int J Infect Dis, 2020. 91: p. 264–266.

7. Li, Y., et al., Strategies for Prevention and Control of the 2019 Novel Coronavirus Disease in the Department of Kidney Transplantation. Transpl Int, 2020.

8. Wu, F., et al., A new coronavirus associated with human respiratory disease in China. Nature, 2020. 579(7798): p. 265–269.

9. Badawi, A. and S.G. Ryoo, Prevalence of comorbidities in the Middle East respiratory syndrome coronavirus (MERS-CoV): a systematic review and meta-analysis. Int J Infect Dis, 2016. 49: p. 129–33.

10. Pallesen, J., et al., Immunogenicity and structures of a rationally designed prefusion MERS-CoV spike antigen. Proc Natl Acad Sci U S A, 2017. 114(35): p. E7348–E7357.

11. Lu, R., et al., Genomic characterisation and epidemiology of 2019 novel coronavirus: implications for virus origins and receptor binding. Lancet, 2020. 395(10224): p. 565–574.

12. Qian, Z.

13. Wang, D., et al., Clinical course and outcome of 107 patients infected with the novel coronavirus, SARS-CoV-2, discharged from two hospitals in Wuhan, China. Crit Care, 2020. 24(1): p. 188.

14. Narayanan, K., et al., Characterization of the coronavirus M protein and nucleocapsid interaction in infected cells. J Virol, 2000. 74(17): p. 8127–34.

15. Zhang, H., et al., Angiotensin-converting enzyme 2 (ACE2) as a SARS-CoV-2 receptor: molecular mechanisms and potential therapeutic target. Intensive Care Med, 2020. 46(4): p. 586–590.

16. Hoffmann, M., et al., SARS-CoV-2 Cell Entry Depends on ACE2 and TMPRSS2 and Is Blocked by a Clinically Proven Protease Inhibitor. Cell, 2020. 181(2): p. 271–280.e8.

17. Nieto-Torres, J.L., et al., Severe acute respiratory syndrome coronavirus envelope protein ion channel activity promotes virus fitness and pathogenesis. PLoS Pathog, 2014. 10(5): p. e1004077.

18. Yang, Y., et al., Traditional Chinese Medicine in the Treatment of Patients Infected with 2019-New Coronavirus (SARS-CoV-2): A Review and Perspective. Int J Biol Sci, 2020. 16(10): p. 1708–1717.

19. Sanders, J.M., et al., Pharmacologic Treatments for Coronavirus Disease 2019 (COVID-19): A Review. Jama, 2020.

20. Pachetti, M., et al., Emerging SARS-CoV-2 mutation hot spots include a novel RNA-dependent-RNA polymerase variant. J Transl Med, 2020. 18(1): p. 179.

21. Kumar, S., et al., MEGA X: Molecular Evolutionary Genetics Analysis across Computing Platforms. Mol Biol Evol, 2018. 35(6): p. 1547–1549.

22. Waterhouse, A.M., et al., Jalview Version 2--a multiple sequence alignment editor and analysis workbench. Bioinformatics, 2009. 25(9): p. 1189–91.

23. Krogh, A., et al., Predicting transmembrane protein topology with a hidden Markov model: application to complete genomes. J Mol Biol, 2001. 305(3): p. 567–80.

24. Apweiler, R., The InterPro database, an integrated documentation resource for protein families, domains and functional sites. Nucleic Acids Research, 2001. 29(1): p. 37–40.

25. Hunter, S., et al., InterPro in 2011: new developments in the family and domain prediction database. Nucleic Acids Res, 2012. 40(Database issue): p. D306–12.

26. Doytchinova, I.A. and D.R. Flower, VaxiJen: a server for prediction of protective antigens, tumour antigens and subunit vaccines. BMC Bioinformatics, 2007. 8: p. 4.

27. Jespersen, M.C., et al., BepiPred-2.0: improving sequence-based B-cell epitope prediction using conformational epitopes. Nucleic Acids Res, 2017. 45(W1): p. W24–W29.

28. Kolaskar, A.S. and P.C. Tongaonkar, A semi-empirical method for prediction of antigenic determinants on protein antigens. FEBS letters, 1990. 276(1-2).

29. Parker, J.M.R., D. Guo, and R.S. Hodges, New Hydrophilicity Scale Derived from High-Performance Liquid Chromatography Peptide Retention Data: Correlation of Predicted Surface Residues with Antigenicity and X-ray-Derived Accessible Sites. Biochemistry, 1986. 25(19): p. 5425–5432.

30. Saha, S. and G.P. Raghava, Prediction of continuous B-cell epitopes in an antigen using recurrent neural network. Proteins, 2006. 65(1): p. 40–8.

31. Bui, H.H., et al., Development of an epitope conservancy analysis tool to facilitate the design of epitope-based diagnostics and vaccines. BMC Bioinformatics, 2007. 8: p. 361.

32. Fleri, W., et al., The Immune Epitope Database and Analysis Resource in Epitope Discovery and Synthetic Vaccine Design. Front Immunol, 2017. 8: p. 278.

33. Vita, R., et al., The immune epitope database (IEDB) 3.0. Nucleic Acids Res, 2015. 43(Database issue): p. D405–12.

34. Zhang, Q., et al., Immune epitope database analysis resource (IEDB-AR). Nucleic Acids Res, 2008. 36(Web Server issue): p. W513–8.

35. Agatonovic-Kustrin, S. and R. Beresford, Basic concepts of artificial neural network (ANN) modeling and its application in pharmaceutical research. Journal of Pharmaceutical and Biomedical Analysis, 2000. 22(5): p. 717–727.

36. Bhasin, M. and G.P. Raghava, Prediction of CTL epitopes using QM, SVM and ANN techniques. Vaccine, 2004. 22(23-24): p. 3195–204.

37. Lamiable, A., et al., PEP-FOLD3: faster de novo structure prediction for linear peptides in solution and in complex. Nucleic Acids Res, 2016. 44(W1): p. W449–54.

38. Dallakyan, S. and A.J. Olson, Small-molecule library screening by docking with PyRx. Methods Mol Biol, 2015. 1263: p. 243–50.

39. Sanner, M.F., Python: A programming language for software integration and development. Journal of Molecular Graphics and Modelling, 1999. 17(1): p. 57–61.

40. Trott, O. and A.J. Olson, AutoDock Vina: improving the speed and accuracy of docking with a new scoring function, efficient optimization, and multithreading. J Comput Chem, 2010. 31(2): p. 455–61.

41. Kelley, L.A. and M.J. Sternberg, Protein structure prediction on the Web: a case study using the Phyre server. Nat Protoc, 2009. 4(3): p. 363–71.

42. Wang, Q., et al., Interaction of α-cyperone with human serum albumin: Determination of the binding site by using Discovery Studio and via spectroscopic methods. Journal of Luminescence, 2015. 164: p. 81–85.

43. Pettersen, E.F., et al., UCSF Chimera--a visualization system for exploratory research and analysis. J Comput Chem, 2004. 25(13): p. 1605–12.

44. Bui, H.H., et al., Predicting population coverage of T-cell epitope-based diagnostics and vaccines. BMC Bioinformatics, 2006. 7: p. 153.

45. De Groot, A.S. and R. Rappuoli, Genome-derived vaccines. Expert Rev Vaccines, 2004. 3(1): p. 59–76.

46. Fauci, A.S., Emerging and re-emerging infectious diseases: influenza as a prototype of the host-pathogen balancing act. Cell, 2006. 124(4): p. 665–70.

47. Korber, B., M. LaBute, and K. Yusim, Immunoinformatics comes of age. PLoS Comput Biol, 2006. 2(6): p. e71.

48. Purcell, A.W., J. McCluskey, and J. Rossjohn, More than one reason to rethink the use of peptides in vaccine design. Nat Rev Drug Discov, 2007. 6(5): p. 404–14.

49. Ahmed, S.F., A.A. Quadeer, and M.R. McKay, Preliminary Identification of Potential Vaccine Targets for the COVID-19 Coronavirus (SARS-CoV-2) Based on SARS-CoV Immunological Studies. Viruses, 2020. 12(3).

50. Lapelosa, M., et al., In silico vaccine design based on molecular simulations of rhinovirus chimeras presenting HIV-1 gp41 epitopes. J Mol Biol, 2009. 385(2): p. 675–91.

51. Chakraborty, S., et al., A computational approach for identification of epitopes in dengue virus envelope protein: a step towards designing a universal dengue vaccine targeting endemic regions. In Silico Biol, 2010. 10(5-6): p. 235–46.

52. Hasan, M.A., M. Hossain, and M.J. Alam, A computational assay to design an epitope-based peptide vaccine against saint louis encephalitis virus. Bioinformatics and Biology Insights, 2013. 7(10): p. 347–355.

53. Frieman, M., M. Heise, and R. Baric, SARS coronavirus and innate immunity. Virus Res, 2008. 133(1): p. 101–12.

54. Pang, J., et al., Potential Rapid Diagnostics, Vaccine and Therapeutics for 2019 Novel Coronavirus (2019-nCoV): A Systematic Review. J Clin Med, 2020. 9(3).

55. Gralinski, L.E. and V.D. Menachery, Return of the Coronavirus: 2019-nCoV. Viruses, 2020. 12(2).

56. Yong, C.Y., et al., Recent Advances in the Vaccine Development Against Middle East Respiratory Syndrome-Coronavirus. Front Microbiol, 2019. 10: p. 1781.

57. Welsh, R.M., L.K. Selin, and E. Szomolanyi-Tsuda, Immunological memory to viral infections. Annu Rev Immunol, 2004. 22: p. 711–43.

58. Shrestha, B. and M.S. Diamond, Role of CD8+ T cells in control of West Nile virus infection. J Virol, 2004. 78(15): p. 8312–21.

59. Maenaka, K. and E.Y. Jones, MHC superfamily structure and the immune system. Current Opinion in Structural Biology, 1999. 9(6): p. 745–753.

60. Reche, P.A. and E.L. Reinherz, Sequence Variability Analysis of Human Class I and Class II MHC Molecules: Functional and Structural Correlates of Amino Acid Polymorphisms. Journal of Molecular Biology, 2003. 331(3): p. 623–641.

61. Sharmin, R. and A.B. Islam, A highly conserved WDYPKCDRA epitope in the RNA directed RNA polymerase of human coronaviruses can be used as epitope-based universal vaccine design. BMC Bioinformatics, 2014. 15: p. 161.

